# Differential efficiency of sampling devices in the measurement of microbial diversity of Yellowstone National Park hot springs

**DOI:** 10.64898/2026.06.15.732322

**Authors:** Jason M. Wood, Scott Tighe, Camilla Urbaniak, Ceth Parker, Nitin Kumar Singh, Season Wong, Brent M. Peyton, Kasthuri Venkateswaran

## Abstract

Metagenomic characterization of low-biomass Yellowstone National Park (YNP) hot spring waters remains challenging because microbial recovery is influenced by filtration methodology, sample preservation, DNA extraction, and sequencing strategy. We characterized thermophilic microbial communities in alkaline YNP hot spring waters (62–90.5°C) using three high-temperature-compatible filtration systems (Sterivex, Supor, and polycarbonate membranes), automated onsite DNA extraction (µTitan), and shotgun metagenomic sequencing with Illumina short-read and Oxford Nanopore Technologies (ONT) long-read platforms. Across all filtration systems and sequencing workflows, microbial communities were consistently dominated by Bacteria (∼90% of reads), whereas Archaea represented <10% of recovered sequences. Dominant microbial populations were reproducibly recovered across all approaches; however, recovery of lower-abundance taxa varied among methods. This variability was most evident in polycarbonate-filtered samples, which exhibited greater replicate-to-replicate variation and less consistent detection of microbial species. *Thermocrinis ruber* and related *Aquificae*-associated thermophiles dominated the hottest waters (78.5–90.5°C), whereas warmer effluent-channel waters (63.5–66.5°C) contained *T. ruber* together with photosynthetic taxa, including *Synechococcus* spp. and *Candidatus Thermochlorobacter aerophilum*. Archaeal communities were primarily represented by *Pyrobaculum*- and *Thermoproteus*-related taxa. Non-metric multidimensional scaling analyses indicated that overall community structure was largely unaffected by filtration or sequencing methodology, whereas alpha-diversity metrics showed that filter selection influenced richness and diversity estimates. These findings identify field-deployable workflows for metagenomic characterization of low-biomass thermophilic aquatic systems and demonstrate the importance of integrating filtration and sequencing strategies for studying extremophile microbiomes under remote sampling conditions.

**IMPORTANCE:** Accurate characterization of low-biomass geothermal water microbiomes remains challenging because microbial recovery is strongly influenced by sample handling, filtration efficiency, DNA extraction chemistry, and sequencing methodology. This study demonstrated that Yellowstone National Park alkaline hot spring water microbiomes were consistently dominated by Bacteria (>90% of recovered reads), whereas Archaea represented <10% of community abundance across all filtration systems. Although dominant microbial populations were reproducibly recovered, filtration-device selection influenced the recovery of microbial diversity and low-abundance taxa. By integrating field-deployable onsite DNA extraction with ONT shotgun metagenomic sequencing, this work evaluates practical workflows for studying thermophilic planktonic microbial communities under remote field conditions. These findings are relevant, not only to geothermal microbiology, but also to low-biomass environments in medical, pharmaceutical, and aerospace industries, where rapid onsite processing and contamination-aware workflows are essential for preserving authentic microbial signatures in extreme environments.

## INTRODUCTION

Studies of Yellowstone National Park (YNP) hot spring ecosystems have provided foundational insights into the ecology, evolution, and functional adaptation of extremophilic microorganisms inhabiting high-temperature geothermal environments (1, 2). Metagenomic investigations across YNP thermal features have revealed extensive microbial diversity associated with variations in temperature, pH, sulfur chemistry, dissolved minerals, and hydrological conditions (3, 4). Accurate characterization of geothermal water microbiomes remains dependent on sample collection, filtration, preservation, and DNA extraction procedures (5, 6). In low-biomass environments, even minor perturbations introduced during sampling and processing can substantially influence downstream metagenomic interpretation (5).

Most previous YNP metagenomic studies focused on microbial mats and sediment-associated communities because these matrices contain substantially higher microbial biomass, which yields sufficient nucleic acids for sequencing with conventional laboratory workflows (2, 6). Microbial mats and sediments frequently contain approximately 10^8^–10^10^ cells/g and support dense microbial consortia structured by steep geochemical gradients. In contrast, planktonic hot spring waters typically contain ∼10^6^ cells/mL, requiring filtration of large water volumes to obtain sufficient material for shotgun metagenomic sequencing (1, 7).

Hot spring water samples also present significant technical challenges for molecular analysis. Most previous YNP water studies relied on workflows in which samples were collected in the field and transported to the laboratory for downstream filtration and DNA extraction (1, 2, 7). Such workflows may introduce physicochemical and biological alterations during transport, cooling, and storage (5). Thermophilic microorganisms adapted to extreme in situ conditions may undergo rapid physiological stress following removal from their native thermal environment, potentially resulting in selective cell lysis, nucleic acid degradation, altered community composition, and reduced recovery of fragile or low-abundance taxa (8-10).

Therefore, immediate onsite processing may improve recovery and preservation of geothermal microbial communities by minimizing transport-associated artifacts and preserving authentic microbial signatures (5, 6, 11). To date, systematic evaluation of field-deployable extraction workflows for YNP geothermal waters has remained limited (6). In addition, elevated temperatures exceeding 80°C in many geothermal systems pose practical challenges for conventional filtration devices and field-based sample handling protocols (1, 8). To address these limitations, a portable onsite processing workflow was evaluated using the validated Microgravity Tested InsTrument for Automated Nucleic Acid Extraction (µTitan) system (5, 6, 11). Following biomass concentration, the µTitan system enabled immediate DNA extraction under remote field conditions, thereby minimizing delays between sample collection and molecular processing (5, 6).

This study evaluated field-deployable workflows for metagenomic characterization of low-biomass thermophilic microbial communities in Yellowstone National Park hot spring waters. Three high-temperature-compatible filtration systems were assessed across a temperature range of 63.5°C to 90.5°C, followed by immediate onsite DNA extraction using the µTitan platform and shotgun metagenomic sequencing on Illumina and Oxford Nanopore Technologies (ONT) platforms. We hypothesized that filtration methodology and sequencing platform would influence recovery of low-abundance taxa but not alter characterization of dominant microbial populations. Because the 2019 dataset consistently yielded >90% bacterial sequences, we evaluated whether onsite filtration and µTitan-based DNA extraction introduced bias against archaeal recovery. To address this question, additional samples collected in 2022 were processed using conventional laboratory-based DNA extraction and amplification-free native nanopore sequencing to independently assess archaeal recovery.

## MATERIALS AND METHODS

### Site description

All field experiments were conducted in YNP at alkaline geothermal features in the White Creek thermal area, located in the Lower Geyser Basin, on June 18, 2019 (Fig. 1). Spring water samples were collected from the Five Sisters Springs (44.5325 °N, 110.7971 °W), Octopus Spring (44.5340 °N, 110.7978 °W), and effluent channels from these features (see Table 1 for a list of samples). The temperature and pH of each site was measured *in situ* using a HQ30d combined pH-temperature probe (Hach, Loveland, CO, USA).

**Table 1.** Yellowstone National Park hot spring samples collected during this study.

| Location | Geothermal Site | Description of the samples collected with | # of samples | pH | T°C |
| --- | --- | --- | --- | --- | --- |
| FSS3 | Five Sisters Spring 3 | Sterivex filter from source pool | 3 | 8.38 | 81.3 |
| FSS3 | Five Sisters Spring 3 | Polycarbonate filter from source pool | 2 | 8.38 | 81.3 |
| FSS3 | Five Sisters Spring 3 | Supor filter from source pool | 2 | 8.38 | 81.3 |
| FSS1 | Five Sisters Spring 1 | Sterivex filter from source pool | 3 | 8.82 | 78.5 |
| FSS1 | Five Sisters Spring 1 | Polycarbonate filter from source pool | 2 | 8.82 | 78.5 |
| FSS1 | Five Sisters Spring 1 | Supor filter from source pool | 2 | 8.82 | 78.5 |
| FSSe2 | FSS effluent channel, closer to creek | Sterivex filter from source pool | 3 | 8.18 | 63.5 |
| FSSe2 | FSS effluent channel, closer to creek | Polycarbonate filter from source pool | 2 | 8.18 | 63.5 |
| FSSe2 | FSS effluent channel, closer to creek | Supor filter from source pool | 2 | 8.18 | 63.5 |
| OSe | OS effluent channel | Sterivex filter from source pool | 3 | 7.94 | 66.5 |
| OSe | OS effluent channel | Polycarbonate filter from source pool | 2 | 7.94 | 66.5 |
| OS | Octopus Spring | Sterivex filter from source pool | 3 | 7.99 | 90.5 |
| OS | Octopus Spring | Polycarbonate filter from source pool | 2 | 7.99 | 90.5 |
| CTL | Control | Sterivex filter control for water (field control) | 3 | NA | NA |
| CTL | Control | Polycarbonate filter control for water (field control) | 1 | NA | NA |
| CTL | Control | Supor filter control for water (field control) | 1 | NA | NA |

Abbreviations: FSS, Five Sisters Springs; OS, Octopus Spring; CTL, control

**Figure 1.**
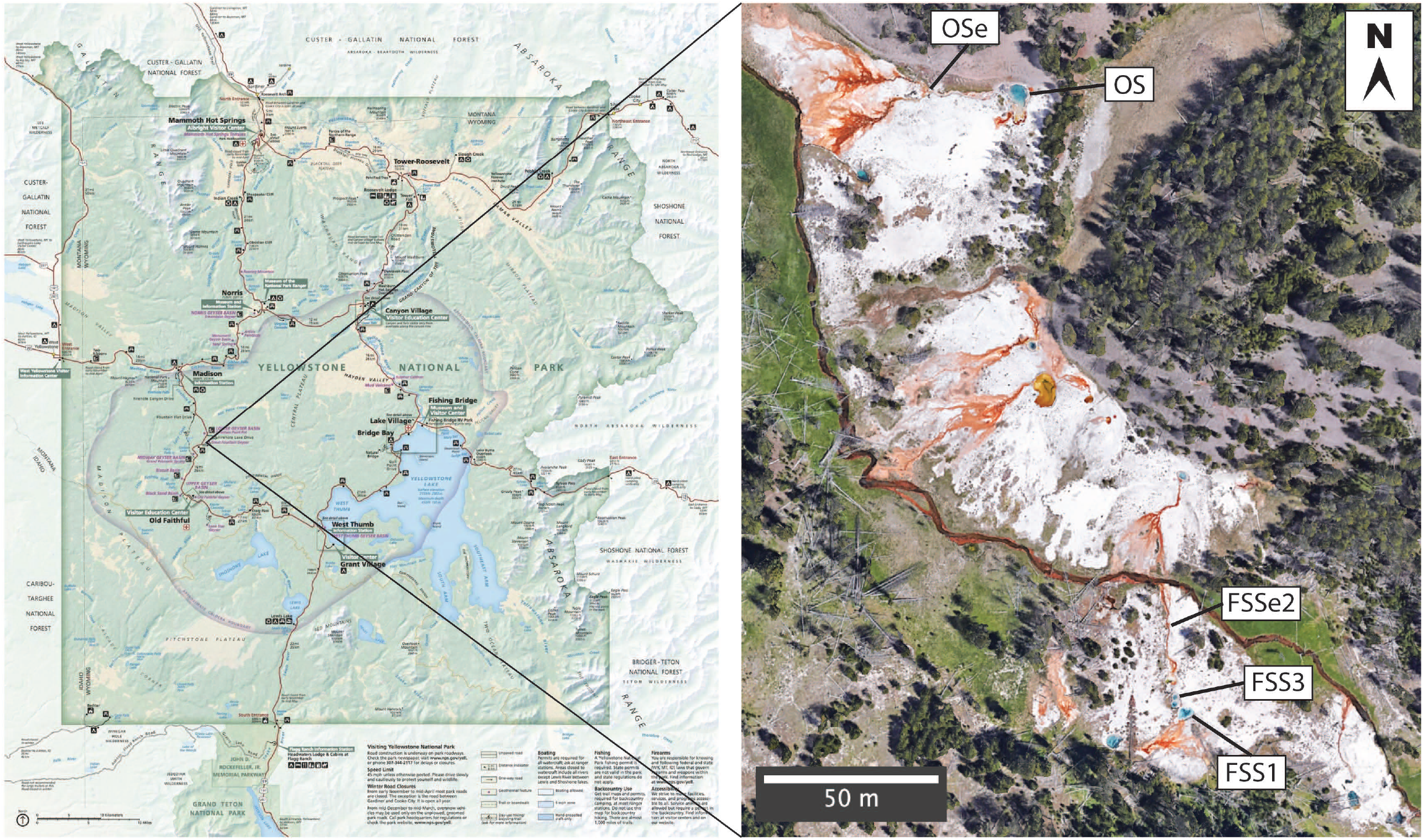
Sampling locations and study design. Geographic locations of Yellowstone National Park hot springs included in this study.

### Physical and chemical parameters of the sampled springs

Vent fluid temperatures ranged between 68°C to 85°C, with the effluent channels of Octopus Spring and Five Sisters Springs measuring between 63.5°C to 90.5°C. Aqueous chemistry was measured from water samples taken either directly from the vent fluid or above the photosynthetic mats. The White Creek thermal area is generally categorized as a water-dominated alkaline-chloride system (12), which is fed from subsurface boiling in deep-heated reservoirs. Collectively, the sampling results all feature high chloride (309 ± 10 mg/L), high arsenic (716 ± 29 µg/L), and low sulphate (18 ± 1 mg/L) concentrations. In addition, the sampled springs are alkaline (pH 7.99–8.85) and contain high concentrations of sodium (140 ± 6 mg/L Na^+^), which is typical of the sodium and chloride-rich waters that form alkaline-chloride hot spring fluids (1).

### On-site DNA extraction using µTitan system workflow

#### (A) Hot spring water sample collection

Prior to sampling, 7.6 m lengths of Masterflex L/S 15 platinum-cured silicone tubing (Cole-Parmer, Vernon Hills, IL, USA) were cut into varying lengths (1.5 m, 3 m, and 3.8 m sections) and sterilized by submerging the tubing into 10% bleach solution for 20 mins. Each tubing section was then rinsed three times with sterile distilled water, coiled, and then wrapped in pre-autoclaved aluminum foil for transport to the field site. In-line polycarbonate 47 mm diameter filter holders (Pall Corporation, Port Washington, NY, USA) were wrapped in aluminum foil and sterilized by autoclave.

To collect water samples, the pump head of a Geopump Peristaltic Pump (GeoTech, Denver, CO, USA) that was equipped with a long section of the sterile tubing was then unwrapped, while keeping both ends remained covered in foil to minimize contamination. Tubing was secured in place, leaving approximately 30 cm on the outflow side of the pump. An inline filter holder was attached to the outflow tubing using a second shorter piece of tubing to the outflow side of the filter holder, with the distal end placed into a graduated cylinder for measurement of filtered volume. The filter holder was subsequently unscrewed and opened. Flame-sterilized tweezers were used to position 47 mm filter membranes into the holder, reassembled, and visually inspected to ensure that no cross-threading, twisting, tearing of the membrane, or gasket displacement had occurred.

The water inflow end of the tubing was lowered in the center of the hot spring using a sampling pole to a location just below the water surface. The pump was initiated and filtration was performed until 10 L was processed or filter clogging occurred. Following the filtration completion, the filter assembly was aseptically disassembled, and the filter was aseptically transferred directly into a screwcap 5 mL tube (VWR, Randor, PA, USA). This procedure was repeated at each site using both 47 mm diameter 0.22 µm Supor filters and 47 mm diameter 0.45 µm polycarbonate filters (Pall Corporation, Port Washington, NY, USA).

Following collection of in-line membrane filter samples, the filter holder was exchanged with an in-line Sterivex-GP Pressure 0.22 µm Filter Unit (Millipore-Sigma, Burlington, MA, USA) using male 3/16th luer barb converters (Cole-Parmer, Vernon Hills, IL, USA) and female 3/16th luer barb converters (Eldon James, Denver, CO, USA). Upon completion of filtration, Sterivex filters were stored in 50 mL Falcon tubes (Thermo Fisher Scientific, Waltham, MA, USA). This was repeated at each sampling location.

#### (B) On-site sample processing for DNA extraction

Each filter was extracted by adding 0.5 mL PBS along with Lysing Matrix E beater beads (MP Biomedicals, Irvine, CA, USA) to the 5 mL tube. Bead beating was performed for 60 s at 20K oscillations per minute using a handheld bead beater tool similar to the BioSpec SoniBeast Jr (JobPLUS ONE 18 V multi tool with P246 console & P570 Head attachment, Ryobi, Anderson, SC, USA). Following bead beating, 25 µL of 10 µg/µL MetaPolyzyme (MAC4L-DF, Millipore Sigma, St. Louis, MO, USA) was added to each tube and incubated for 1 hr at 37°C, followed by another round of bead beating for 15 s. Tubes were then allowed to settle for 15 min, before aliquoting 100 µL of the supernatant into the µTitan system for DNA extraction.

#### (C) On-site DNA extraction

A commercially available magnetic particle–based nucleic acid isolation kit (NucliSENS Magnetic Particle Extraction Kit, bioMerieux, Durham, NC, USA) used as the chemistry in the µTitan DNA extraction system (11). Each sample was extracted in triplicate and simultaneous negative controls were processed using molecular-grade water (Sigma-Aldrich, St. Louis, MO, USA) and processed alongside the samples.

### Sequencing workflow

#### (A) Samples processed in 2019

Laboratory-based Oxford Nanopore sequencing was performed using the GridION-X5 sequencer equipped with MinION Rev 9.4.1 flow cells. Libraries were synthesized using the rapid PCR Barcoding kit (SQK-RPB004) with 3 µL of adjusted input volume. PCR cycles were adjusted for total input amounts as follows; 16 cycles for 1 to 3 ng, 18 cycles for >0.25 to 1 ng, and 22 cycles for <0.25 ng. Read depth varied for each sample, and low reads were obtained for samples that had DNA at or below detection limits according to the results of the Qubit dsDNA HS Assay kit (ThermoFisher Scientific, Waltham MA). While low read depth was noted for some samples that had high inputs, likely reflecting low microbial biomass, biofilm-associated interference, or the presence of inhibitory compounds. Whole genome shotgun library synthesis was performed on 1ng input or maximum input volume using the Nextera XT whole genome kit (Illumina Inc., San Diego, CA, USA), cleaned, quantified using the Qubit spectrofluorometer (Thermo Fisher Scientific, Waltham, MA, USA), and analyzed for quality using the Bioanalyzer 2100 (Agilent Technologies, Santa Clara, CA, USA). Libraries were equimolar pooled and sequenced using single-end 150 bp chemistry on a HiSeq 1500/2500 system (Illumina, San Diego, CA, USA). Samples were demultiplexed, and fastq files were used for analysis.

#### (B) Samples processed in 2022

To independently assess whether the predominance of bacterial sequences (>90%) observed in the 2019 dataset was influenced by the µTitan-based DNA extraction workflow, additional water samples were collected from Five Sisters Springs in September 2022 and analyzed using an established laboratory workflow (7). In contrast to the 2019 samples, which were field-filtered and subjected to immediate onsite DNA extraction, the 2022 samples were transported on ice, filtered in the laboratory, and processed using a conventional DNA extraction method (7) followed by amplification-free native nanopore sequencing. Briefly, one-liter samples were collected and filtered using a 47mm 0.22 µm MCE membrane (Millipore Cat. GSWP04700) and DNA extracted using the DNeasy PowerWater kit (Qiagen Cat. 14900). DNA was quantified using Qubit dsDNA HS Assay kit followed by native nanopore direct DNA sequencing using 100 ng of gDNA following the manufacturer recommended protocol for the Native barcoding kit (Oxford Nanopore NDB114-24). Samples were sequenced by pooling 20 fmol of each barcoded library and loading a Rev10.4 P2 flow cell (PRO114M) until more than 2x10^6^ reads were obtained for each sample.

### Sequence quality control and annotation

Quality control of short-read Illumina sequence data was performed using Trimmomatic (v0.32) to remove adapter sequences and low-quality reads, using a quality cutoff value set at minimum Phred score of 20 along the length of the read. Quality control of Nanopore sequence data was performed using Nanofilt (v2.6.0) and default parameters. Reads shorter than 80 bp were removed from the data. The DIAMOND aligner (v2.0.4) was used to query the NCBI-NR protein database, and the MEGAN 6 lowest common ancestor (LCA) algorithm was used to bin high quality reads to their respective taxonomy. Additionally, the eggnog, KEGG, and SEED databases were queried for functional analysis. For long-read Nanopore sequence annotation, the DIAMOND frameshift and range curling mode was used. The frameshift option performs frameshift-alignment of DNA sequences against a protein reference database. The parameter used with the frameshift mode was -F 15, leading to a dynamic programming penalty of 15. Range-culling estimates an alignment locally; this option was used with the parameter as follows --range-culling and --top 10.

Taxonomic classification, relative abundance determination, and sequence quality analyses of native nanopore sequence datasets generated in 2022 samples were conducted using the One Codex platform (One Codex, Wilmington, DE, USA) with default parameters. Sequence reads were assigned taxonomic identities using the One Codex k-mer–based classification algorithm against a curated microbial reference database (13, 14).

### Statistical Analyses

To determine whether long-read ONT sequencing could supplement or replace short-read Illumina sequencing for more rapid analyses in remote locations, multiple statistical analyses were performed. Alpha diversity among similar samples was measured with Chao1, Shannon, and Simpson indices as calculated by the R vegan package (15), with rarefication just under the size of the smallest library (N=5000). The R vegan package (15) was also used to visualize beta diversity using nonmetric multidimensional scaling (NMDS) of Bray-Curtis dissimilarity of rarefied samples.

## RESULTS AND DISCUSSION

Results from this study included water samples collected at the Five Sister and Octopus Springs sites at Yellowstone National Park on June 18, 2019. A detailed description, temperature, and pH of each of the samples are given in Table 1. YNP hot spring water samples were collected from five different locations using Sterivex, Supor, and polycarbonate filtration systems and characterized by comprehensive microbiome analyses employing multiple whole-genome sequencing approaches, including Illumina and Oxford Nanopore technologies (Fig. 2). Replicate samples (n=2 and 3) were collected at each site to characterize the reproducibility of the µTitan DNA extraction workflow and sequencing platforms. In total, 31 water samples (15 Sterivex, 10 polycarbonate, and 6 Supor filters) and five field controls (3 Sterivex, 1 polycarbonate, and 1 Supor) were included in the analysis (Table 1). Sequencing read counts for each detected species were combined across replicate samples collected from the same location prior to calculating relative abundance. To minimize background noise and improve consistency among datasets, a minimum threshold of 10 reads per taxa was used for ONT datasets, and >500 reads for Illumina datasets which resulted in maintaining >99.9% of total reads.

**Figure 2.**
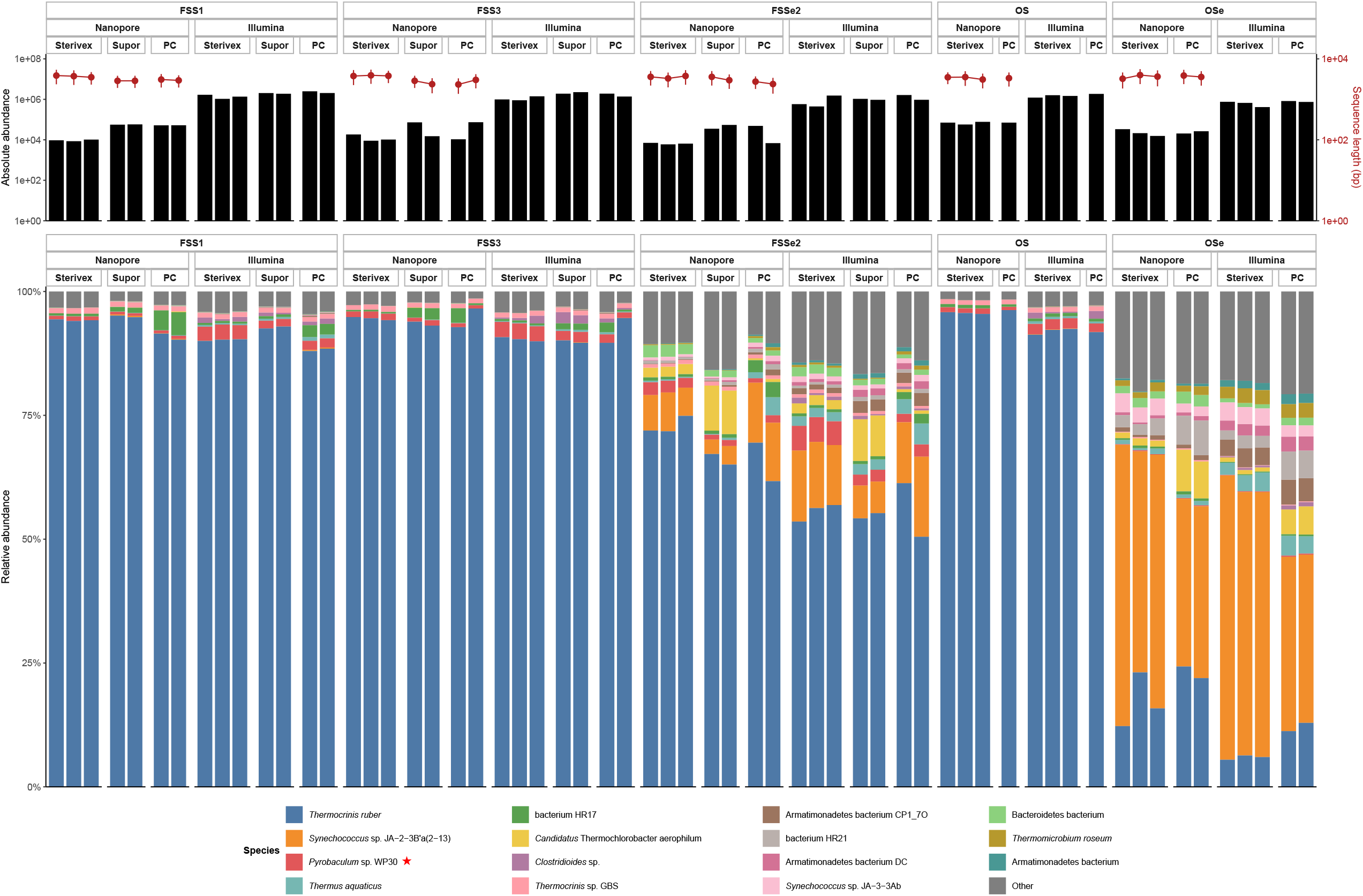
Effects of filtration method and sequencing platform on microbial abundance and community composition. Comparison of microbial abundance, sequencing performance, and taxonomic composition among Yellowstone hot spring samples processed using Sterivex, Supor, and polycarbonate (PC) filtration devices and analyzed by Illumina and Oxford Nanopore sequencing. Upper panels show absolute microbial abundance estimates (black bars) and sequence read lengths (red symbols). Lower panels depict species-level relative abundances of dominant bacterial and archaeal (red asterisk) taxa recovered from each sample.

### Archaeal diversity

Archaeal community composition recovered by the different filtration systems is presented (Table 2). When Illumina sequencing was used, the Sterivex filtration system recovered more archaeal reads across all Five Sisters Spring water samples, with the greatest recovery observed in the hotter Five Sisters Spring (81.3°C; FSS3) and Octopus Spring (90.5°C; OS) samples. Similar to the total reads retrieved, higher numbers of archaeal taxa were recovered when the Sterivex filtration system was used (Table 2). In general, ∼62% to 70% of *Pyrobaculum* sp. WP30 sequences were recovered by all three filtration systems in hot water (81.3°C and 78.5°C) samples of Five Sisters Spring and Octopus Spring (90.5°C) samples. The archaeal taxa not recovered by the polycarbonate and Supor filters represented <3% of total reads and did not alter the interpretation of dominant community composition.

**Table 2.** Dominant archaeal populations found in water samples from YNP springs using all three filtration systems.

| Method | Archaeal taxa | Five Sisters Spring 3 (81.3°C) |  |  | Five Sisters Spring 1 (78.5°C) |  |  | FSS effluent 2 (63.5°C) |  |  | Octopus Spring (90.5°C) |  | OS effluent (66.5°C) |  |
| --- | --- | --- | --- | --- | --- | --- | --- | --- | --- | --- | --- | --- | --- | --- |
|  |  | FSS3 |  |  | FSS1 |  |  | FSSe2 |  |  | OS |  | Ose |  |
|  |  | Sterivex | PC | Supor | Sterivex | PC | Supor | Sterivex | PC | Supor | Sterivex | PC | Sterivex | PC |
| Illumina shotgun short reads: |  |  |  |  |  |  |  |  |  |  |  |  |  |  |
|  | *Pyrobaculum* sp. WP30 | 62.1 | 67.3 | 65.9 | 64.1 | 64.9 | 66.8 | 55.4 | 65.6 | 66.5 | 70.3 | 74.2 | 55.1 | 55.4 |
|  | *Pyrobaculum* sp. OCT_11 | 7.0 | 7.5 | 6.9 | 5.7 | 5.9 | 6.2 | 6.4 | 7.2 | 7.4 | 8.2 | 8.4 |  |  |
|  | *Thermoproteus* sp. CP80 | 6.6 | 6.1 | 6.6 | 6.6 | 7.0 | 7.1 | 6.3 | 7.0 | 7.0 | 4.9 | 5.3 |  |  |
|  | *Pyrobaculum* sp. JCHS_4 | 5.1 | 5.1 | 5.0 | 5.3 | 5.2 | 5.4 | 4.7 | 5.1 | 5.3 | 4.7 | 5.2 |  |  |
|  | archaeon HR02 | 0.7 | 1.1 | 0.8 | 0.7 | 1.5 | 1.3 | 1.0 | 1.9 | 1.1 | 0.6 | 0.6 | 10.1 |  |
|  | Candidatus *Caldiarchaeum subterraneum* |  | 0.3 | 0.3 | 0.3 | 0.6 |  | 0.8 | 1.9 | 0.8 |  | 0.2 | 34.8 | 34.0 |
|  | archaeon HR01 | 0.5 |  |  |  |  |  |  |  |  |  |  |  | 10.6 |
|  | Others | 18.1 | 12.6 | 14.6 | 17.3 | 14.8 | 13.3 | 25.4 | 11.3 | 11.9 | 11.3 | 6.0 |  |  |
| # of taxa |  | 25 | 15 | 20 | 25 | 20 | 17 | 41 | 14 | 14 | 17 | 9 | 3 | 3 |
| Total # of reads |  | 164,968 | 71,738 | 128,930 | 184,647 | 130,899 | 91,467 | 226,154 | 79,211 | 69,362 | 128,851 | 42,998 | 5,155 | 7,657 |
| ONT long reads: |  |  |  |  |  |  |  |  |  |  |  |  |  |  |
|  | *Pyrobaculum* sp. WP30 | 59.1 | 76.4 | 67.6 | 59.5 | 65.7 | 69.3 | 53.3 | 68.0 | 58.0 | 71.9 | 78.1 | 19.3 | 23.3 |
|  | *Pyrobaculum* sp. OCT_11 |  | 7.3 | 5.7 |  | 4.7 | 5.3 |  | 3.4 | 4.4 | 6.4 | 7.0 | 2.1 |  |
|  | *Thermoproteus* sp. CP80 | 7.2 | 4.3 | 6.1 | 8.7 | 6.6 | 5.3 | 9.5 | 6.1 | 5.8 | 4.6 | 5.1 | 2.1 |  |
|  | archaeon HR02 |  |  |  |  | 4.3 | 2.8 |  | 4.1 | 1.6 | 1.2 |  | 4.7 | 4.2 |
|  | Candidatus *Caldiarchaeum subterraneum* |  |  |  |  | 2.3 | 1.5 |  | 5.2 | 1.3 |  |  | 48.2 | 52.6 |
|  | Thermoprotei archaeon | 5.9 |  | 3.0 | 9.0 | 5.0 | 4.1 | 5.5 | 2.5 | 4.2 |  |  |  | 3.4 |
|  | Desulfurococcaceae archaeon AG1 | 5.9 | 1.5 | 1.8 | 5.9 | 1.8 | 2.1 | 5.8 |  | 3.4 |  |  |  |  |
|  | archaeon HR01 |  |  |  |  |  |  |  | 1.2 |  |  |  | 9.2 | 7.0 |
|  | Euryarchaeota archaeon |  |  |  |  |  |  |  |  | 1.1 |  |  | 4.2 | 5.7 |
|  | Others | 22.0 | 10.5 | 15.9 | 16.9 | 9.6 | 9.6 | 25.9 | 9.6 | 20.2 | 15.9 | 9.8 | 10.3 | 3.9 |
| # of taxa |  | 13 | 8 | 15 | 9 | 12 | 11 | 13 | 10 | 25 | 19 | 6 | 12 | 7 |
| Total # of reads |  | 801 | 740 | 1,104 | 390 | 950 | 921 | 860 | 815 | 1,785 | 2,944 | 590 | 622 | 386 |

When ONT sequencing was used, the Supor filtration system recovered more high molecular weight DNA fragments compared to the Sterivex system for Five Sisters Springs samples (Table 2) and had notable improvements to the polycarbonate workflow, both on generating reads and archaeal taxa for all hot springs samples tested during this study. Unlike Illumina, where all filtration systems invariably recovered higher proportion of sequences of *Pyrobaculum* sp. WP30, only <60% of the total archaeal reads were assigned to *Pyrobaculum* sp. WP30 except for Octopus Spring (90.5°C) samples (∼72%). As observed with Illumina sequencing, the nanopore platform also recovered dominant archaea (*Pyrobaculum* sp.) from water samples filtered through all three systems.

### Bacterial diversity

Differential measurement of bacterial population using various filtration systems as measured by all samples being extracted using the µTitan platform and sequenced using Illumina and Oxford nanopore technology platforms for various hot springs water samples (n=16; Table 3). Unlike archaea, more bacterial diversity was observed from the Supor filters when the Illumina system was used for hot (81.3°C) samples of Five Sisters Spring water. The Supor filtration system yielded both higher total read recovery and greater bacterial taxonomic richness in the hot (81.3°C) Five Sisters Spring water samples. The bacterial taxa not recovered by the Sterivex workflow represented approximately 4% of total reads and did not affect interpretation of the dominant community composition. Relative abundance of *Thermocrinis ruber* sequences were observed to be ∼93% to 96% in all three filtration systems in hot water (81.3°C and 78.5°C) samples of Five Sisters Spring and Octopus Spring (90.5°C). In the warmer Five Sisters Spring sample (66.3°C), *T. ruber* sequences accounted for only ∼58–63% of reads, with the highest abundance recovered by the Sterivex system for the Five Sister Springs warm (66.3°C) samples, whereas Supor (58%) had somewhat similar but lower incidence. Conversely, the polycarbonate system retrieved more bacteria (69 taxa) in the Octopus Spring hot (90.5°C) samples when compared to the Sterivex system. In addition, when Octopus Spring warm water (66.5°C) was filtered through the polycarbonate filtration system, *Synechococcus* sp. sequences were only 0.03% of total reads whereas Sterivex was >56%. The polycarbonate filter overwhelmingly retrieved *T. ruber* sequences (>95%) but only 6% recovered with the Sterivex workflow. Furthermore, Sterivex retrieved more bacterial populations (60 taxa) while only 25 taxa were observed with polycarbonate filtered Octopus Spring warm (66.5°C) water samples.

**Table 3.** Dominant bacterial populations found in water samples from YNP springs using all three filtration systems.

| Method | Bacterial taxa | Five Sisters Spring 3 (81.3°C) |  |  | Five Sisters Spring 1 (78.5°C) |  |  | FSS effluent 2 (63.5°C) |  |  | Octopus Spring (90.5°C) |  | OS effluent (66.5°C) |  |
| --- | --- | --- | --- | --- | --- | --- | --- | --- | --- | --- | --- | --- | --- | --- |
|  |  | FSS3 |  |  | FSS1 |  |  | FSSe2 |  |  | OS |  | OSe |  |
|  |  | Sterivex | PC | Supor | Sterivex | PC | Supor | Sterivex | PC | Supor | Sterivex | PC | Sterivex | PC |
| Illumina shotgun short reads: |  |  |  |  |  |  |  |  |  |  |  |  |  |  |
|  | *Thermocrinis ruber* | 96.0 | 94.6 | 93.7 | 95.7 | 92.0 | 95.8 | 63.0 | 60.4 | 58.0 | 95.5 | 12.5 | 6.1 | 95.1 |
|  | *Thermocrinis* sp. GBS | 1.0 | 0.9 | 0.9 | 1.0 | 0.9 | 1.0 | 0.8 | 0.6 | 0.6 | 0.9 | 0.1 | 0.1 |  |
|  | *Synechococcus* sp. JA-2-3B'a(2-13) |  |  |  | 0.1 |  |  | 14.4 | 14.4 | 6.9 | 0.1 | 35.9 | 56.3 | 0.0 |
|  | Armatimonadetes bacterium CP1_70 |  |  |  |  |  |  | 1.3 | 2.4 | 2.5 | 0.1 | 5.1 |  |  |
|  | *Thermus aquaticus* | 0.2 | 0.3 | 0.4 | 0.3 | 0.8 | 0.4 | 2.1 | 3.6 | 2.2 |  |  | 3.1 | 0.4 |
|  | *Synechococcus* sp. JA-3-3Ab |  |  |  | 0.1 | 0.2 |  | 1.1 | 1.1 | 0.9 |  | 2.3 | 3.6 |  |
|  | Bacterioidetes bacterium |  |  |  | 0.1 | 0.1 |  | 2.0 | 0.9 | 1.2 |  | 1.6 | 0.9 |  |
|  | Others |  |  |  |  |  |  |  |  |  |  |  |  |  |
| # of taxa |  | 32 | 42 | 43 | 39 | 51 | 39 | 76 | 79 | 68 | 45 | 69 | 60 | 25 |
| Total # of reads |  | 3,107,450 | 3,191,080 | 4,006,357 | 3,864,193 | 4,370,059 | 3,810,570 | 2,277,435 | 2,450,173 | 1,886,166 | 4,132,385 | 1,514,273 | 1,791,870 | 1,794,828 |
| ONT long reads: |  |  |  |  |  |  |  |  |  |  |  |  |  |  |
|  | *Thermocrinis ruber* | 97.7 | 97.4 | 95.5 | 97.6 | 92.7 | 96.5 | 79.1 | 70.7 | 68.3 | 97.3 | 97.5 | 16.8 | 23.6 |
|  | *Thermocrinis* sp. GBS | 1.1 | 0.9 | 1.0 | 1.2 | 1.0 | 1.0 | 0.8 |  |  | 1.0 | 1.0 | 0.2 |  |
|  | *Synechococcus* sp. JA-2-3B'a(2-13) |  |  |  |  |  |  | 7.5 | 12.4 | 3.5 |  |  | 53.3 | 35.3 |
|  | *Synechococcus* sp. JA-3-3Ab |  |  |  |  |  |  | 0.8 | 0.8 | 0.8 |  |  | 3.5 | 1.9 |
|  | Bacterioidetes bacterium |  |  |  | 0.1 |  |  | 2.5 | 1.0 | 1.4 |  |  | 1.7 | 2.4 |
|  | Candidatus *Thermochlorobacter aerophilum* |  |  | 0.1 |  | 0.1 | 0.1 | 2.2 | 0.4 | 9.2 |  |  | 1.3 | 8.1 |
|  | Others |  |  |  |  |  |  |  |  |  |  |  |  |  |
| # of taxa |  | 16 | 24 | 40 | 12 | 50 | 42 | 42 | 74 | 94 | 35 | 23 | 89 | 88 |
| Total # of reads |  | 36,500 | 83,491 | 86,282 | 27,262 | 102,180 | 112,167 | 18,005 | 53,801 | 86,668 | 201,694 | 69,828 | 68,545 | 45,707 |

The ONT sequencing demonstrated greater levels of high molecular weight DNA for bacteria using the Supor filtration system when compared to the Sterivex system for Five Sisters Spring samples (Table 3) while the polycarbonate system recovered greater bacterial diversity when compared to Sterivex for Five Sisters Spring samples. Sterivex filtration nevertheless yielded higher recovery of overall reads and taxa from Octopus Spring samples. As seen in Illumina short read sequences, all filtration systems invariably recovered a higher proportion of sequences of *T. ruber* (>95%) for all hot samples (>78.5°C); whereas the relative abundance of *T. ruber* was lower in the warmer samples of Five Sisters Springs (∼68 to 79%) and Octopus Spring (∼16% to 24%). As documented by short-read sequencing, the ONT platform also recovered dominant bacteria (>95%) when water samples were filtered through all three systems, with the remaining reads comprising taxa at a relative abundance of 0.5% to 1%.

The consistent dominance of *Thermocrinis/Aquificae*-associated bacterial lineages across filtration systems and sequencing workflows indicates that the major community members were robustly recovered regardless of analytical approach. Previous studies of the YNP geothermal ecosystems frequently emphasized archaeal dominance, particularly within acidic sediments and microbial mat-associated communities where anaerobic microenvironments, sulfur cycling, and steep thermal gradients promote archaeal proliferation (1, 2). The present study, however, showed that planktonic alkaline hot spring waters may support relatively similar bacterial-dominated communities despite the presence of thermophilic archaea. Although these findings differ from previous 16S-based observations (7), the methods and sample types used are different and differences in microbial diversity are not unexpected. *Thermocrinis* species are well known hydrogen- and sulfur-oxidizing thermophiles that thrive in oxygenated alkaline geothermal waters (16) and are commonly associated with *Aquificae*-dominated spring outflows and mixing zones (17, 18).

### Alternate sequencing approach to support microbial diversity profile

Selected hot spring water samples collected in 2022 were analyzed using amplification-free native nanopore sequencing following conventional laboratory-based DNA extraction to independently characterize microbial community composition. This approach provided a ground-truth assessment of community structure by eliminating biases associated with PCR amplification and library preparation. Sequencing quality metrics demonstrated successful recovery of high-molecular-weight environmental DNA. For sample FSS1, read lengths ranged from approximately 0.5 to 42 kb with an N50 of 7,532 bp, whereas sample FSS3 yielded reads ranging from 0.5 to 41 kb with an N50 of 5,826 bp. These results indicate that the extracted DNA was of sufficient quality to support long-read sequencing and taxonomic characterization.

The taxonomic profiles obtained from the 2022 samples were consistent with those recovered in the 2019 dataset. Native nanopore sequencing recovered microbial communities composed of approximately 90% Bacteria and >10% Archaea (Fig. 3), whereas the corresponding Rapid PCR Barcoding (SQK-RPB004) workflow recovered approximately 95% Bacteria and ∼5% Archaea (data not shown), suggesting improved recovery of archaeal sequences with the amplification-free approach. Despite differences in relative abundance estimates, both amplification-based and amplification-free workflows identified the same dominant bacterial and archaeal populations, supporting the conclusion that thermophilic alkaline hot spring waters were predominantly inhabited by bacterial communities. To our knowledge, this study represents the first application of native nanopore sequencing for characterization of Yellowstone hot spring water microbiomes and highlights the value of amplification-free long-read sequencing for generating robust and minimally biased assessments of microbial diversity in extreme environments.

**Figure 3.**
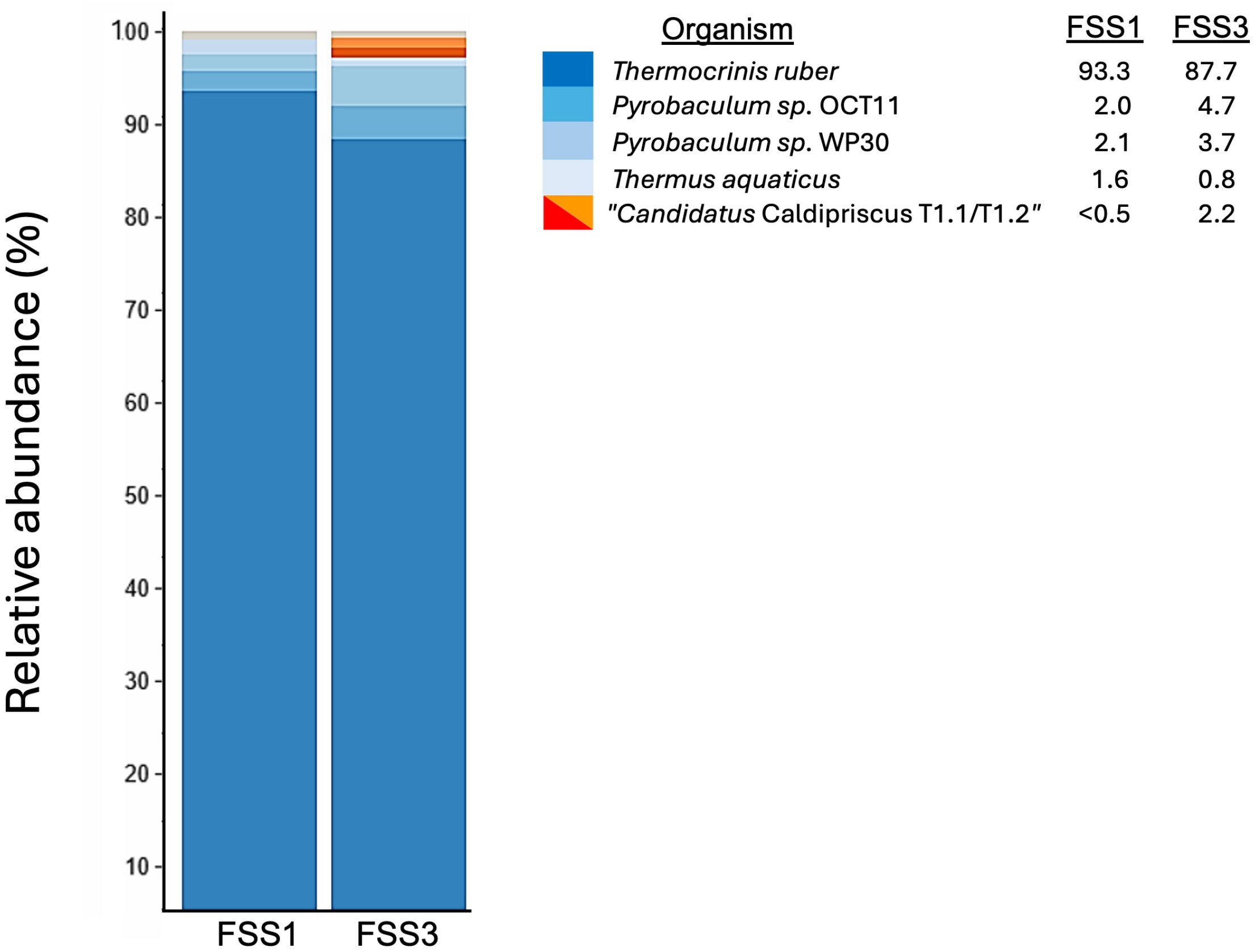
Native nanopore sequencing validates microbial community composition. Species-level taxonomic composition of Yellowstone National Park hot spring water samples (FSS1 and FSS3) determined using amplification-free native nanopore sequencing.

### Influence of filtration material on microbial recovery

Many previous YNP studies have used polycarbonate filters (19) to concentrate microorganisms from liquid samples, since they are rated to withstand higher temperatures during filtration (∼140°C). In order to confirm the differential filtration of microbes from hot water, we tested multiple filtration systems and found that Sterivex filtration system detected more taxa using short read sequencing (Illumina) of archaea in all Five Sister Spring water samples, especially from hot samples of both Five Sisters Spring (81.3°C; FSS3) and one of the Octopus Spring (90.5°C; OS) samples. One explanation is that short-read sequencing provides greater sequencing depth and enables detection of shorter DNA possibly including fragmented eDNA. Nanopore sequencing revealed that the Supor filtration recovered greater archaeal taxa when compared to Sterivex system for samples from Five Sisters Springs, possibly because nanopore is less likely to sequence shorter fragmented DNA from poorly intact organisms. Sterivex filtration also outperformed the polycarbonate system on recovering archaeal taxa for all hot springs samples tested during this study. Likewise, the Sterivex workflow detected a greater diversity of bacteria (60 taxa) while only 25 taxa were observed from the polycarbonate method for Octopus Spring (66.5°C) water samples. Of note is that at higher temperatures, the outer plastic of the Sterivex flow through filter could be seen to warp, although this did not affect recovery of Archaea or Bacterial taxa.

### Sequencing platforms

Differences between short-read sequencing (Illumina-based) and nanopore sequencing (Oxford Nanopore) workflows were evident throughout the study. Illumina sequencing generated substantially higher read depth and more consistent recovery of low-abundance taxa, resulting in tight clustering of replicate samples in NMDS analyses (Fig 4) and greater reproducibility in alpha diversity (Fig 5) measurements. ONT sequencing produced longer reads (3 to 8 kb) for rapid PCR barcoding (RPB004) and 0.5 to 41 kb for the native DNA sequencing (NDB114) approach which improved taxonomic continuity and enabled long-read mapping across thermophilic genomes, although total read depth remained lower and variability among replicate samples was greater. This variability likely reflects stochastic sampling effects and library-preparation biases associated with low-biomass samples. Similar sequencing-depth limitations associated with ONT workflows have been reported in other low-biomass environmental metagenomic studies (9, 10). Despite these differences, dominant taxa were consistently recovered by both platforms.

**Figure 4.**
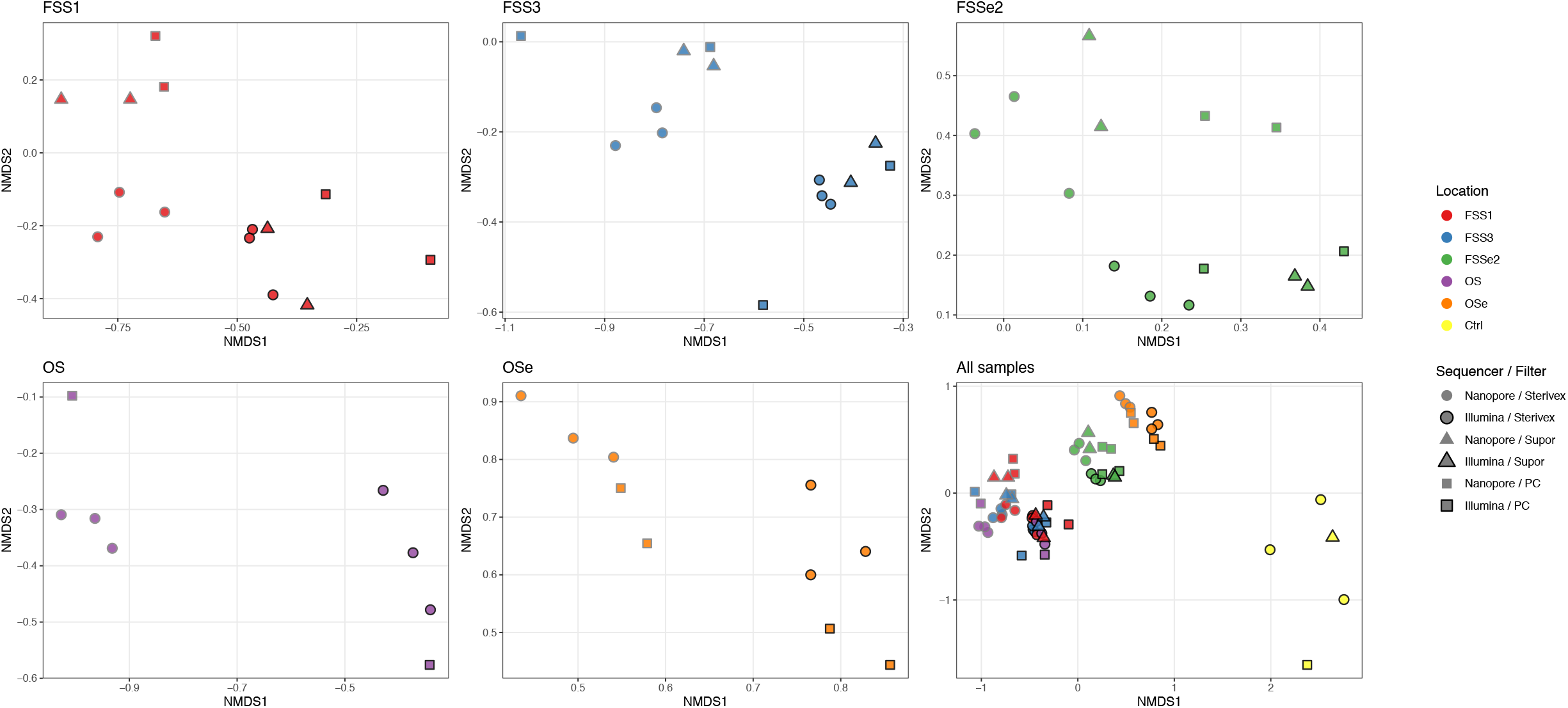
NMDS analysis of microbial community structure. Ordinations are based on Bray– Curtis dissimilarity matrices showing the effects of sampling location, filtration device, and sequencing platform on microbial community composition. Individual panels represent separate hot spring locations (FSS1, FSS3, FSSe2, OS, and OSe), while the combined ordination illustrates relationships among all samples including negative controls. Colors denote sampling locations, and symbols indicate filtration device and sequencing platform combinations. Samples showed limited separation by filtration method or sequencing platform, indicating that methodological effects were secondary to the dominant microbial community structure.

**Figure 5.**
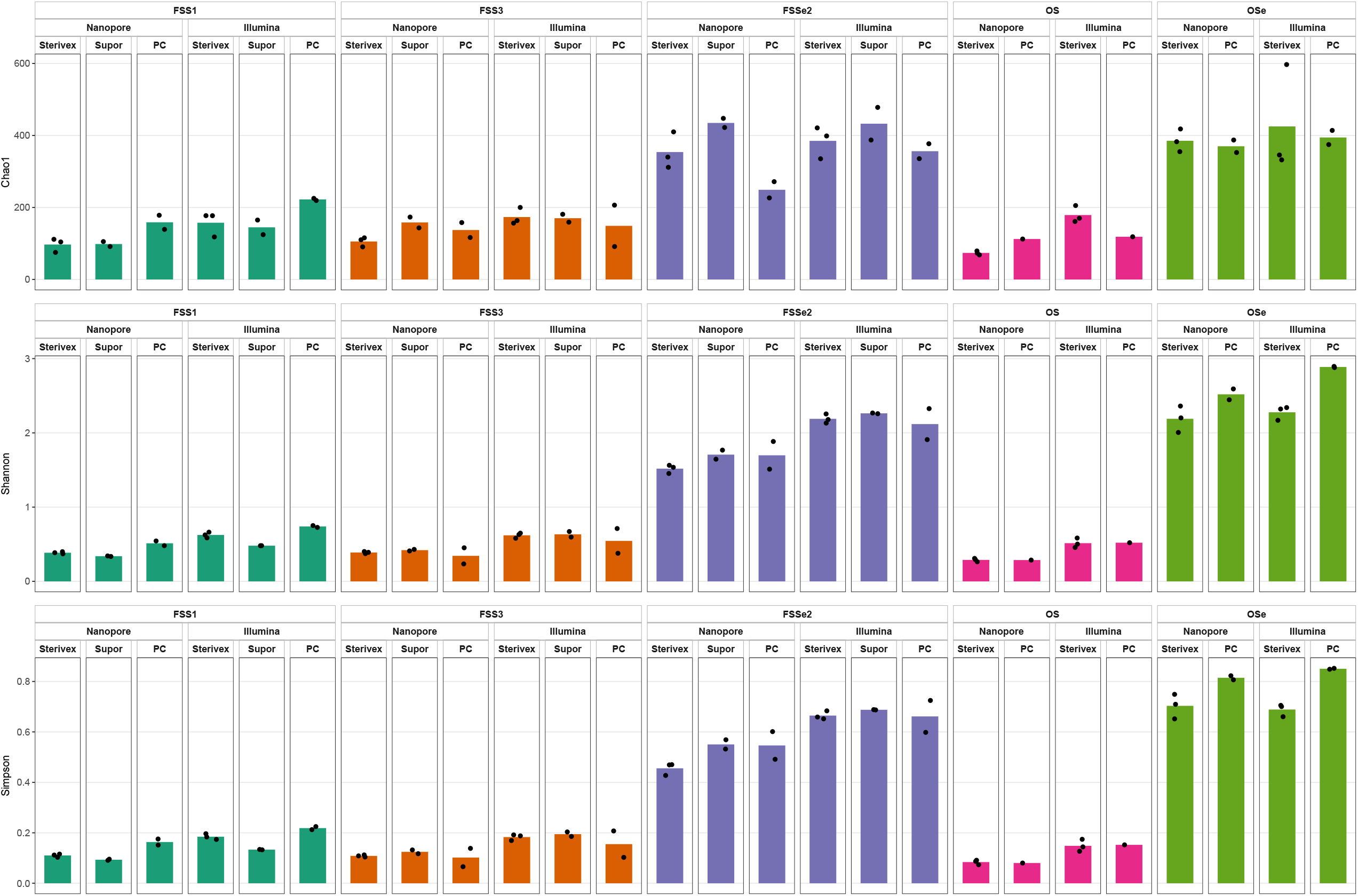
Alpha diversity metrics across filtration devices and sequencing platforms. Alpha diversity estimates for Yellowstone National Park hot spring microbial communities recovered using Sterivex, Supor, and polycarbonate (PC) filtration devices and analyzed by Illumina and Oxford Nanopore sequencing. Richness (Chao1), diversity (Shannon), and evenness (Simpson) indices are shown for each sampling site. Data points represent biological replicates.

### NMDS and community structure analyses

The NMDS analyses further supported the conclusion that characterization of dominant microbial communities remained stable across filtration systems and sequencing approaches (Fig. 4). Biological samples clustered distinctly from field controls, indicating minimal influence of contamination or kit-associated background DNA on overall microbial community structure. Control samples formed separate outlier clusters, particularly in ONT datasets, suggesting that background contaminants contributed minimally to dominant thermophilic microbial signatures. Although subtle separation among filtration systems was observed, particularly within Illumina datasets, the overall clustering patterns demonstrated that filtration methodology exerted secondary effects relative to the strong ecological dominance of shared thermophilic bacterial populations. No strong segregation of samples by hot spring location was observed within the ordination space, indicating that differences among planktonic microbial communities were less pronounced than expected across these closely related alkaline geothermal environments.

### Alpha diversity indices

Alpha diversity analyses indicated that filtration-device selection influenced microbial richness and community evenness (Fig. 5). Polycarbonate filters often yielded higher Chao1 richness and Shannon diversity estimates, particularly in ONT datasets, suggesting improved recovery of rare or low-abundance taxa. These filters have been widely used in geothermal studies because of their high thermal tolerance and maximum operating temperature of 140°C (19). They also exhibited greater variability among replicates, indicating that inconsistencies may arise during downstream processing. In several high-temperature samples, polycarbonate filters recovered lower microbial diversity and reduced abundance of dominant taxa compared with Sterivex and Supor filters. Because repeated microscopic examination of autoclaved polycarbonate filters revealed no evidence of structural damage, the observed variability is unlikely to result from filter degradation. Instead, it may reflect inefficiencies associated with biomass recovery during the extraction and grinding procedures.

Sterivex filters generally provided the most consistent performance across samples, although their use for hot water samples (>65°C) should be approached cautiously because prolonged exposure to elevated temperatures may compromise filter integrity. These findings suggest that filtration performance in extreme geothermal environments depends not only on thermal compatibility but also on the efficiency and reproducibility of downstream sample-processing workflows.

Despite differences in richness estimates, Shannon and Simpson diversity values remained relatively low across all filtration systems and sequencing workflows. These low diversity values reflect the strong predominance of a limited number of thermophilic taxa, particularly *Thermocrinis/Aquificae*-associated bacterial lineages, with only minor contributions from lower-abundance bacterial and archaeal populations. Similar low-evenness communities have been reported in other geothermal systems where environmental extremes impose strong selective pressures favoring specialized thermophilic populations (2, 3). Although filtration methodology influenced recovery of rare taxa and alpha-diversity estimates, the dominant thermophilic community structure remained highly consistent across workflows.

The successful implementation of field-deployable molecular workflow(s) for hot spring waters (∼90°C) represents an important step in the metagenomic characterization of low-biomass geothermal systems under remote sampling conditions. Immediate onsite filtration and portable nucleic acid extraction with a chemistry-adjustable (tunable) platform such as µTitan minimizes the delay between sample collection, filter processing, and molecular processing, thereby reducing opportunities for transport-associated perturbations, selective cell lysis, and nucleic acid degradation (5, 6). Because thermophilic microorganisms adapted to extreme geothermal environments may undergo rapid physiological stress following removal from their native habitat, rapid onsite processing may improve preservation of native microbial signatures and enhance recovery of low-abundance extremophiles (8).

In this study, we evaluated several workflows that were integrated with a field-deployable nucleic acid extraction device. These methods compared large-volume onsite sample concentration (>5 L) followed by real-time DNA extraction with preselected chemistry and platform-matched sequencing. Supor filters are recommended as one of the primary methods when high-molecular-weight DNA and ONT sequencing are priorities, while Sterivex filters provide a complementary option for consistent recovery of dominant bacterial and archaeal taxa but must be used with caution for high temperature samples. Historically, polycarbonate filters have been used for many years (20) and have shown higher capture richness and rare taxa, but have greater replication variability due to potential processing and handling issues.

The workflows evaluated here may also be applicable to other low-biomass extremophile systems relevant to astrobiology (21), planetary protection (22), and remote environmental monitoring applications (5). High-temperature geothermal systems are frequently considered analog environments for early Earth ecosystems and potentially habitable extraterrestrial environments (23-25). These findings provide a framework for developing optimized field-deployable workflows for low-biomass thermophilic microbiomes and other extreme environments relevant to astrobiology and planetary protection, where rapid onsite processing and contamination control are essential (26).

## ACKNOWLEDGEMENTS

This study was conducted under Yellowstone National Park research permit YELL-2019-SCI-5480 held by BMP. We appreciate Dana Skorupa for assistance in the field sampling and challenging discussions. We acknowledge Fathi Karouia and the students and staff members of the Induced Environments Group (formerly Biotechnology and Planetary Protection group) for their continuous technical support, when necessary. © 2026 California Institute of Technology, Government sponsorship acknowledged.

## AUTHORS’ CONTRIBUTIONS

JMW, CU, CP, NKS, SW, BMP, ST, and KV were involved in planning the field expedition to Yellowstone National Park and performed various functions on-site, including sampling, sample processing, and nucleic acid extraction using the µTitan. JMW and BMP provided local knowledge of the sampling site and its dangers. JMW and CP were trained to handle any emergency situations while on-site. BMP contributed to the geochemistry components of this work. ST performed DNA library synthesis, DNA quality control, DNA sequencing using the ONT and Illumina platforms.

JMW and NKS performed bioinformatic analyses. BMP, ST, and KV supervised all work. JMW, ST, and KV contributed to writing this manuscript. All authors have read the manuscript and agree with its content.

## DATA AVAILABILITY

The datasets presented in this study can be found in online repositories. The names of the repository/repositories and accession number(s) can be found at NCBI PRJNA935810.

## DISCLAIMER

This manuscript was prepared as an account of work sponsored by NASA, an agency of the US Government. The US Government, NASA, California Institute of Technology, Jet Propulsion Laboratory, and their employees make no warranty, expressed or implied, or assume any liability or responsibility for the accuracy, completeness, or usefulness of information, apparatus, product, or process disclosed in this manuscript, or represents that its use would not infringe upon privately held rights. The use of, and references to any commercial product, process, or service does not necessarily constitute or imply endorsement, recommendation, or favoring by the US Government, NASA, California Institute of Technology, or Jet Propulsion Laboratory. Views and opinions presented herein by the authors of this manuscript do not necessarily reflect those of the US Government, NASA, California Institute of Technology, or Jet Propulsion Laboratory, and shall not be used for advertisements or product endorsements.

